# Activity of a three-phage combination against *Mycobacterium tuberculosis* in disease-relevant conditions

**DOI:** 10.64898/2026.05.11.724220

**Authors:** S. Janssen, S.E. Larsen, M. Pozuelo Torres, M. Beldjenna, C.A. Guerrero Bustamante, I. Florian, T. Smytheman, T. Guo, R.C. van Wijk, G.F. Hatfull, A.H. Diacon, R.N. Coler, J. van Ingen

## Abstract

Phage therapy offers promise to combat antimicrobial resistance, including drug-resistant tuberculosis (TB). Understanding phage activity against *Mycobacterium tuberculosis* (Mtb) adapted to physiologic microenvironments, such as hypoxia and acidity in granulomas, is essential since these conditions induce non-replicating states. We evaluated a phage combination against Mtb under hypoxic, acidic (pH 5.5), and stationary-phase conditions *in vitro*.

In planktonic *Mtb* growth conditions, phage concentrations increased around day seven followed by a significant reduction in *Mtb* H37Rv load, which was maintained over 31 days. Phage addition prevented regrowth was observed with rifampicin and isoniazid alone. Individual phage stability was differentially affected by acidic media conditions, resulting in variability of antimycobacterial activity. In hypoxic conditions and stationary growth experiments, phage titers remained stable over time with no change in mycobacterial load compared to controls. Model-based predictions were able to adequately capture phage-mycobacterial interactions with and without rifampicin.

The lack of antimycobacterial activity in assays with non-replicating mycobacteria suggest that phages need actively replicating mycobacteria to exert lytic activity. Stable phage concentrations in assays with non-replicating mycobacteria suggests low grade phage replication in these conditions. Established models can support future study design through simulations of different experimental scenarios.

## Introduction

Tuberculosis (TB) remains a major threat to global health(1). With over ten million cases, of which ∼390,000 are diagnosed with rifampicin (RIF-) resistant or multidrug resistant (MDR-) TB annually, emerging resistance and transmission of antibiotic-resistant strains is a major concern(2). Significant progress has been made over the past years with the introduction of an all-oral, 6-9 months regimens for MDR-TB(3). Rising resistance rates to bedaquline, introduced as recently as the early 2010s and now the backbone of this regimen, are already reported(4–6). Alternative or adjunctive approaches to treatment of TB will be required to address this threat.

The increase in antimicrobial resistance (AMR) has fuelled interest in bacteriophage (phage) therapy(7). Unlike antibiotics, phages often have limited host ranges, resulting in the need for individualized treatment targeted at the bacterial strain causative of an infection. However, the genotypic diversity of *M. tuberculosis* (*Mtb*) is relatively low(8), making this bacterium an ideal candidate for development of a standardized multiphage treatment that can be used in a majority of TB patients(9). Recently, a combination of mycobacteriophages with *in vitro* lytic activity against the major human-adapted lineages of *Mtb* complex was developed(9).

*Mtb* can adapt to different physiological conditions encountered within granulomas including hypoxia, acidic environments and intracellular niches within macrophages. This can lead to *Mtb* entering a non-replicating, dormant state making it difficult to kill (10, 11). Such *Mtb* are tolerant to many clinically used antibiotics and result in the required extended duration of antibiotic regimens to treat TB(10). As phages use their bacterial host’s machinery for replication preceding lytic activity, it is unclear whether phages are able to infect and lyse dormant *Mtb* in these conditions.

Understanding the kinetics of phage-mycobacterial interactions in diverse physiologic environments can help define optimal timing of phage treatment and potential antibiotic combinations, particularly if these complex interactions can be predicted by statistical models.

We selected three mycobacteriophages with broad lytic activity across *Mtb* lineages, favourable manufacturing characteristics and sufficient diversity to minimize the risk of *Mtb* phage cross-resistance(9). We investigated the antimycobacterial activity of these phages against *Mtb* H37Rv in a range of physiologic conditions, including planktonic growth, acidic conditions, hypoxia and stationary phase, and investigated if statistical models were able to predict the findings.

## Results

An overview of the general experimental set-up for experiments testing activity of three phages (Fred313cpm-1Δ33 (Fred), FionnbharthΔ45Δ47 (Fionnbharth), Muddy HRM N0157-2 (Muddy)) against *Mtb* H37Rv in normal, acidic (pH 5.5), hypoxic and stationary phase conditions is shown in Figure 1.

**Figure 1.**
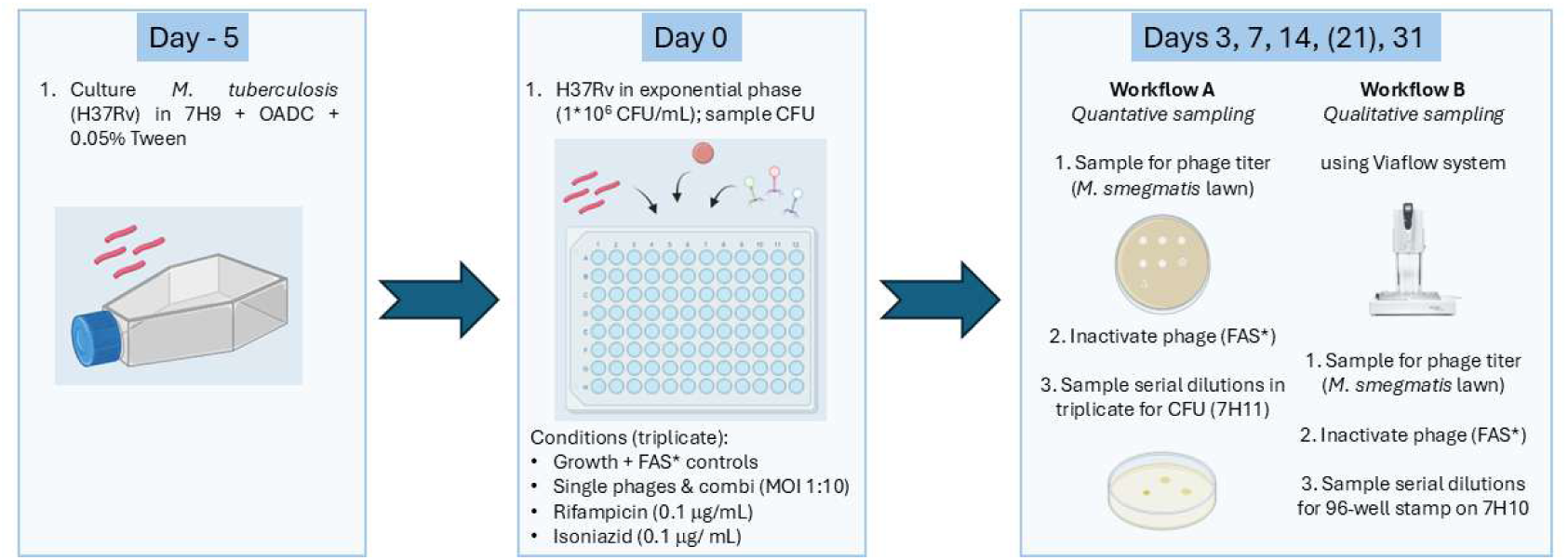
Overview of experimental set-up. *An overview of the general experimental set-up is shown. Abbreviations: colony-forming units (CFU), ferrous ammonium sulphate (FAS), multiplicity of infection (MOI)*.

### Phage activity against replicating M. tuberculosis

In planktonic growth conditions, a reduction in mycobacterial load was observed in all phage-treated conditions compared to the ferrous ammonium sulphide (FAS; phage inactivation buffer) treated growth control from day 10 (Fred, Fionnbharth and phage combination) or day 14 (Muddy) (Figure 2A-B, Supplementary Table). The bactericidal effect of the single phages and the phage combination was sustained over time through the last sampling point at day 31. The bactericidal effect was preceded by an increase in phage concentrations up from day seven (Figure 2C). Phage concentrations were significantly higher for Fionnbharth and Muddy compared to the phage combination, whereas concentrations for Fred were lower. Evaluating plaque titers on phage-specific *M. smegmatis* mutants showed an even representation of all three phages in the phage combination throughout the assay (Supplementary Figure 1-2).

**Figure 2.**
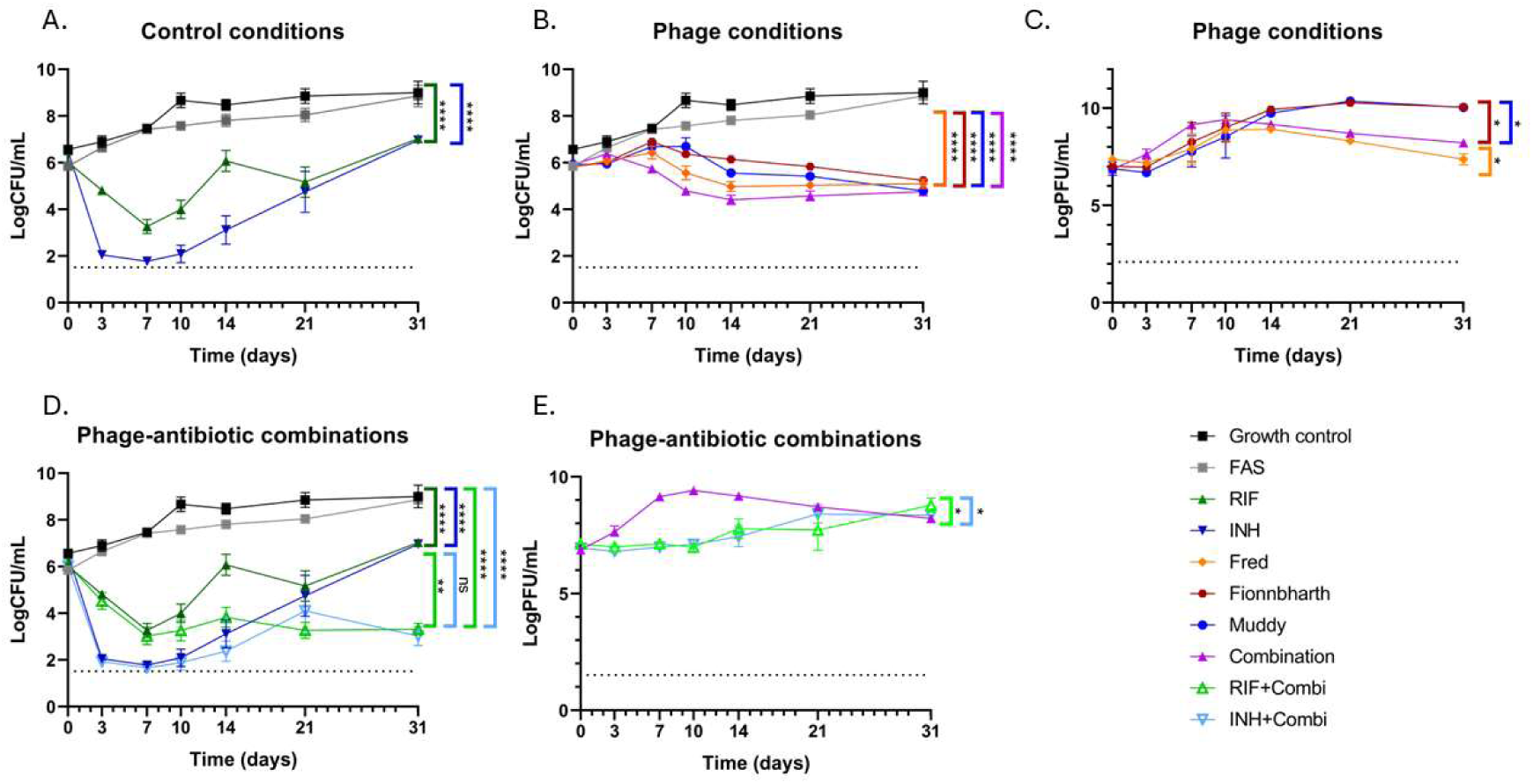
Time kill assay with replicating mycobacteria. *Mean and standard errors are shown for mycobacterial loads (A, B, D)) and phage concentrations (C and E) resulting from time-kill assays with M. tuberculosis and phage. P-values are shown for largest differences between experimental conditions compared to the growth control, or phage-antibiotic combinations versus antibiotics only, respectively. Level of significance: * p<0.05 and ** p<0.01, ***p<0.001, ****p<0.0001. Abbreviations: colony forming units (CFU), plaque forming units (PFU), ferrous ammonium sulphate (FAS), rifampicin (RIF), isoniazid (INH), not significant (ns)*.

A 2-log increase was observed over time in growth control conditions. FAS treatment led to a mild decrease in colony forming units (CFU) with a significantly lower area under the curve (AUC) compared to the growth control without FAS treatment (Figure 2A, Supplementary Table).

With the rifampicin (RIF) and isoniazid (INH) exposure, regrowth of mycobacteria was observed up from day 14 (Figure 2D). This regrowth was prevented when RIF or INH conditions were also treated with the phage combination (Figure 2D). Phage concentrations were higher for the phage combination condition compared to the phage combination with antibiotics, where later increases in phage titers were observed. Phage titers on phage-specific *M. smegmatis* mutants showed an even representation of all three phages in the phage combination together with RIF throughout the assay (Supplementary Figure 2). Isolates harvested on day 31 from the RIF only condition did not demonstrate any mutations associated with resistance, whereas isolates harvested on Day 31 from the INH conditions with and without phage demonstrated mutations on the katG gene (p.Trp328Arg (INH) and p.Glu709Lys (INH+phage), not described as known mutations associated with resistance according to the WHO catalogue(12). The minimum inhibitory concentration was >2 mg/L for both isolates qualifying as resistant to INH.

### Phage activity in acidic conditions

In a time-kill assay with *Mtb* in acidic conditions (pH 5.5), incubation with Muddy and the phage combination led to a decrease in mycobacterial load at day 31 only (Figure 3A-B, Supplementary Table 1), but not to the same magnitude as observed in normal pH media (Figure 2B). There were no differences in mycobacterial load between the other phage treated conditions and the growth controls. A 1-log increase in CFU/mL in growth controls was observed over time. RIF had a significant and sustained bactericidal effect, confirming the assay controls were in alignment with expectations (Figure 3A). Phage concentrations rapidly decreased, except for Muddy and the phage combination in alignment with our phage stability findings without *Mtb* (Figure 3C; Supplementary Figure 3). Evaluating plaque titers on phage-specific *M. smegmatis* mutants suggested the effect in the phage combination was likely driven by Muddy in one series of experiments in acidic media (Supplementary Figure 2). However, these differences were not statistically significant due to inter-experiment variation.

**Figure 3.**
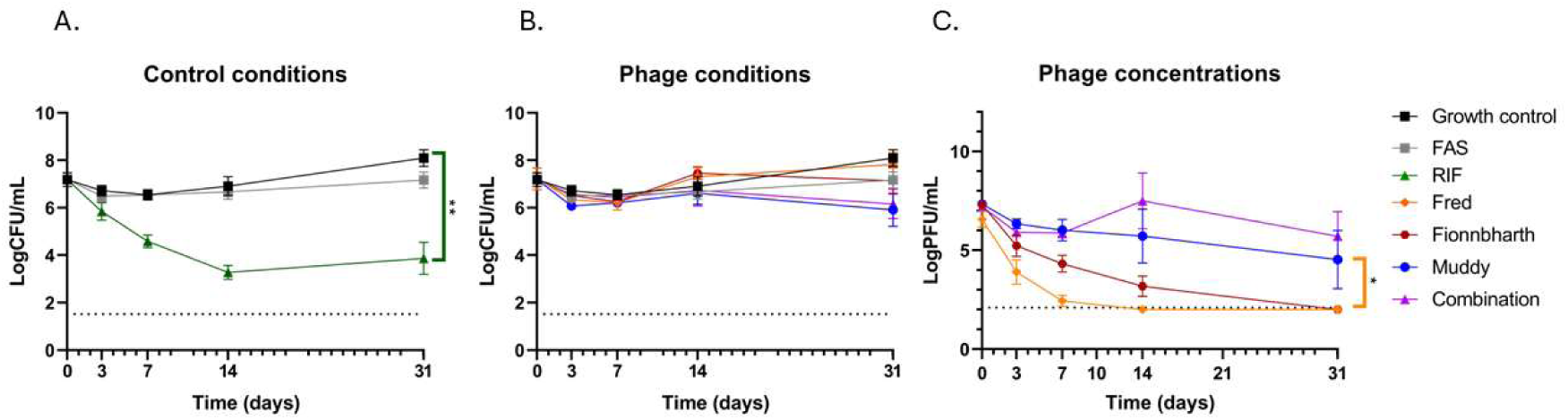
Phage activity in acidic conditions. *Mean and standard errors are shown for mycobacterial loads (A and B) and phage concentrations (C) resulting from time-kill assays with M. tuberculosis and phage at pH 5.5. Level of significance: ****p<0.0001. Abbreviations: colony forming units (CFU), rifampicin (RIF), plaque forming units (PFU), ferrous ammonium sulphate (FAS)*.

### Phage activity against stationary phase M. tuberculosis

Phage added to stationary phase cultures of *Mtb* did not change the bacterial load and these cultures remained stable over time for growth controls and phage treated conditions, except for the phage combination showing a modest effect (Figure 4A-B, Supplementary Table). In this assay, no reduction in mycobacterial load was observed for the RIF control. In the parallel time-kill experiment, bactericidal activity of RIF and the phage combination was as expected (Figure 4C). Phage concentrations in the stationary phase experiment remained stable until day seven, after which a reduction was observed for Fred and Fionnbharth at day 14, whereas Muddy and the phage combination remained stable (Figure 4D).

**Figure 4.**
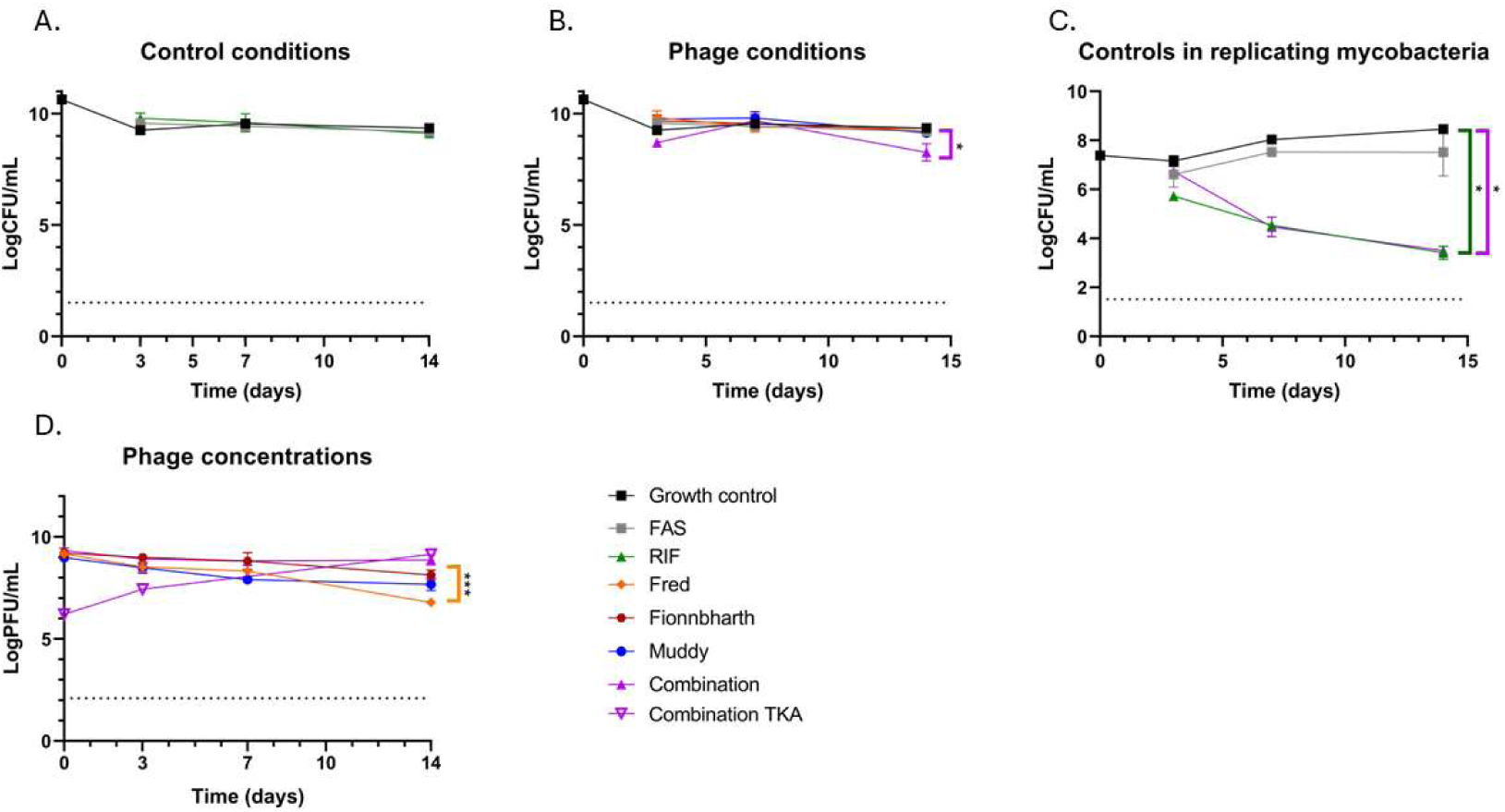
Lack of phage activity against stationary phase mycobacteria. *Mean and standard errors are shown for mycobacterial loads (A and B) and phage concentrations (D) resulting from experiments with stationary phase M. tuberculosis and phage. The results of a parallel time-kill experiment with added fresh medium are also shown (C, D) Level of significance: ****p<0.0001. Abbreviations: colony forming units (CFU), rifampicin (RIF), plaque forming units (PFU), ferrous ammonium sulphate (FAS)*.

### Phage activity against M. tuberculosis at low oxygen tension

The last condition evaluated was hypoxia, as *Mtb* can persist in an array of oxygen gradients and understanding how this may affect phage susceptibility will be key to further applications. In anaerobic conditions the mycobacterial load in the growth controls remained stable over time (Figure 5A). Mycobacterial loads in phage treated conditions also remained stable over time (Figure 5B). Phage concentrations also remained stable over time, except for Fred where a decrease was observed from day seven (three days after adding phage) (Figure 5C). A significant decrease in mycobacterial load was observed for INH and RIF (Figure 5A). The bactericidal effect of both antibiotics was reduced compared to effects observed in time-kill assays with replicating mycobacteria.

**Figure 5.**
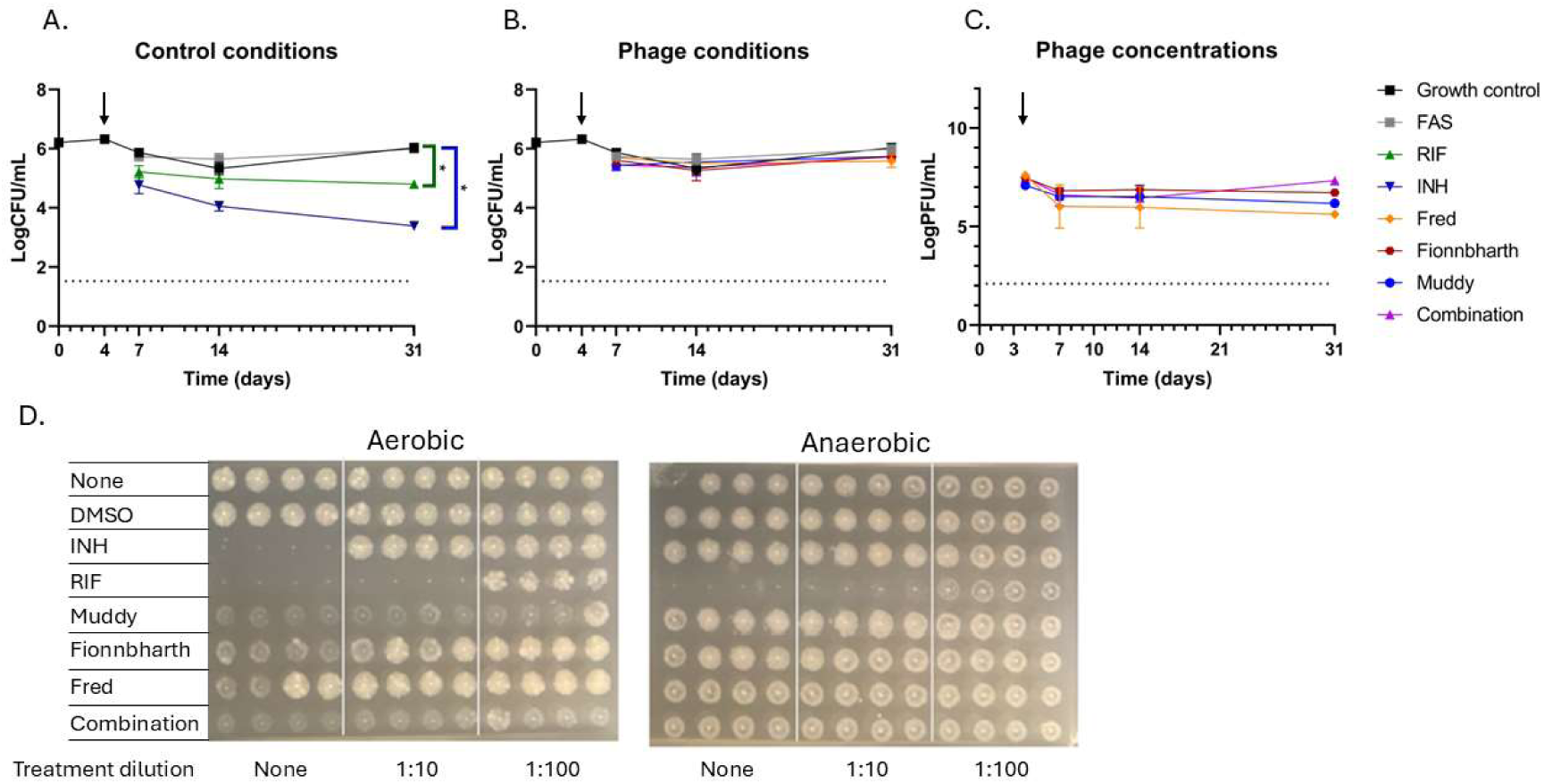
Phage activity in hypoxic conditions. *Mean and standard errors are shown for mycobacterial loads (A and B) and phage concentrations (C) resulting from hypoxia experiments. Arrows indicate the addition of experimental treatment to mycobacteria after adaptation to hypoxia. Omitted wash steps at Day 4 in one replicate of the hypoxia experiment led to a peak in CFU, which was not observed in a replacement experiment with wash steps. CFU from the replacement experiment were used for the final analysis. Panel D shows images of CFU stamps taken with high throughput methodology for parallel aerobic and anaerobic conditions at Day 8 (3 days after addition of treatment; rows show technical triplicates for three dilutions of treatment added)*. *Level of significance: * p<0.05 and ** p<0.01. Abbreviations: colony forming units (CFU), rifampicin (RIF), isoniazid (INH), plaque forming units (PFU), ferrous ammonium sulphate (FAS)*.

To assess whether an established phage infection during hypoxia would lead to a bactericidal effect, in one series at day 31 a plate was returned to oxygen-rich conditions after inactivation of extracellular phage. An increase in mycobacterial load was observed in the growth control and the phage combination treated conditions in aerobic conditions, whereas the mycobacterial load in the anaerobic conditions remained the same (Supplementary Figure 4).

As a complement to the quantitative CFU/mL data, we used a high throughput stamping methodology to generate qualitative data regarding changes in phage effects in hypoxic conditions. On day eight, (day three post treatment), we observed a drop in efficacy for INH, but no change with RIF treatment on CFU spots observed, confirming cultures were in a hypoxia induced state (Figure 5D). We also confirmed a titratable effect of INH and RIF in aerobic conditions. None of the individual phages nor the combination were able to reduce *Mtb* CFU in hypoxic conditions, whereas the effect in regular oxygen conditions was evident for Fionnbharth, Muddy and the phage combination (Figure 5D).

### Phage stability in different conditions

At 4℃, phage concentrations remained stable over time in supplemented 7H9 at pH 6.6 and pH 5.5 conditions until Day 31 with a mild decrease for Fred at pH 5.5. Phage stability was impaired at 37℃ with significant decreases of all phage concentrations in medium with pH 5.5 and decreases for Fred in medium with pH 6.6 (Supplementary Figure 3).

### Model-based predictions of phage-mycobacterial dynamics

To characterize the longitudinal dynamics of *Mtb* and phages, a theoretical model was developed based on the phage combination and RIF conditions in the time kill experiment with replicating mycobacteria. Natural growth of *Mtb* was quantified, and efficacy of RIF (decrease in objective function value [dOFV] = 205.69 with degrees of freedom [df] = 2, p < 0.0001) and phage combination (dOFV = 213.68 with df = 7, p < 0.0001) were significant improvements. Combined RIF with phage combination was predicted based on monotherapy and overlaid for validation (Supplementary Figure 6). Sustained mycobacterial reduction predicted in the phage combination condition corresponded to a preceding increase in PFU (Figure 6A) as captured by the compartmental model structure from phage infection of *Mtb* to its burst (Supplementary Table 2-3). Consistent with the data, the model predicted approximately 2-log increase in control conditions, and regrowth between 1-2 weeks was predicted based on decreased apparent efficacy of RIF. Phage concentrations were lower when combined with RIF, increasing later (Figure 6A).

**Figure 6.**
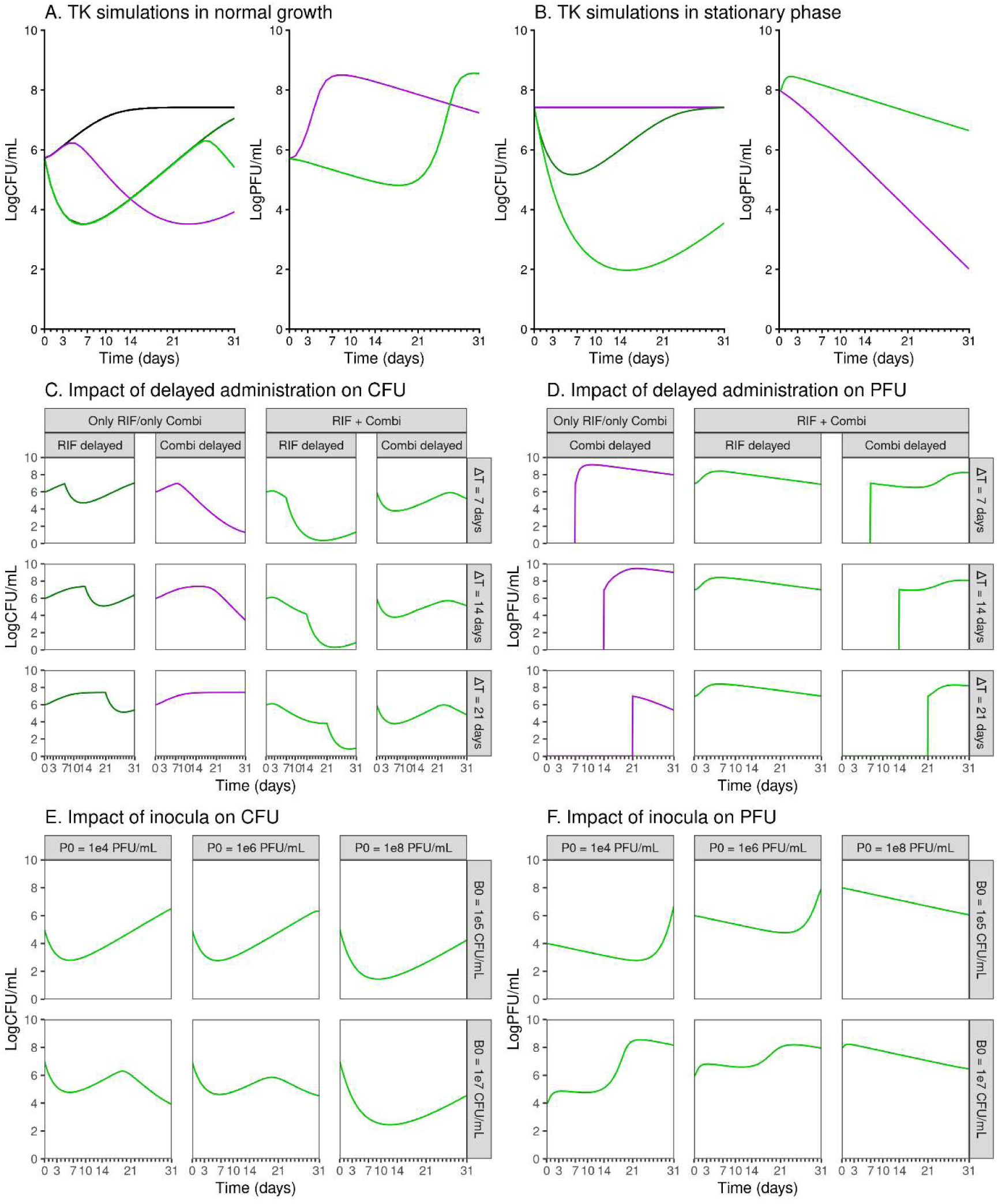
Model simulations of time kill dynamics of mycobacterial and phage load for different scenarios of rifampicin and phage treatment. *Model simulations of time kill dynamics of mycobacterial (log CFU/mL) and phage (log PFU/mL) load for different scenarios of RIF and phage combination treatment. (A) Experimental groups as tested in the replicating Mtb experiment. (B) Bacterial burden and (C) phage load for different baseline bacterial burdens (rows) and phage combination inocula (columns). (D) Bacterial burden and (E) phage load for different delays in time of treatment starts (rows) for antibiotic, combination, or antibiotic plus combination (columns)*. *Abbreviations: B0 = baseline bacterial burden, FAS = ferrous ammonium sulphide, GC = growth control, P0 = phage combination inoculum, RIF = rifampin, TK = time kill, ΔT = delay in time of treatment start*.

The impact of treatment may be optimized, given the interaction between phage, bacterium (predator-prey) and antibiotic. Scenarios of different starting points are simulated, showing a benefit in delaying RIF treatment after phage combination treatment even after 7 days, while delaying phage combination treatment did not benefit efficacy (Figure 6C-D).

When simulating different scenarios of bacterial burden (rows) and different phage inocula (panels), an increased phage efficacy for higher phage inocula can be seen with increased initial decrease in CFU (Figure 6E-F). Interestingly, a higher bacterial burden of 10^7^ CFU/mL shows increased phage efficacy to approximately 4-log decrease at the highest phage inoculum. In addition, a clear predator-prey pattern was characterized. Despite being developed based on replicating *Mtb* data, the model can predict stationary *Mtb* scenarios (Figure 6B) with the phage combination treatment showing no change in *Mtb* bacterial burden and phage load declining over time.

## Discussion

To understand the potential role of phage therapy in future regimens for TB treatment, it is helpful to get insights in the lytic activity of phages against *Mtb* in diverse physiologic conditions, simulating disease-relevant environments within granulomas and macrophages. The activity of the phages was confirmed by the significant and sustained decrease in mycobacterial load after incubation with the single phages and the phage combination in time-kill assays with replicating mycobacteria up from day seven, preceded by an increase in phage concentrations. In predictive models, phages combined with antibiotics have increased bactericidal activity compared to only phages or only antibiotics, even without incorporation of an interaction term. This is in line with the different modes of action of phage and antibiotic activity.

In the disease relevant physiological states, phages proved less effective. The lower and variable phage activity in acidic environments may be related to the decrease in phage concentrations due to instability at pH 5.5 (see Supplementary Figure 2) possibly preventing phages from establishing a replicative infection, or the lower metabolic activity and multiplication rate of the mycobacteria (1 log10 increase in CFU/mL in the growth control versus 2 log increase in planktonic conditions) influencing phage-bacterial interactions. Variability in phage stability in acidic conditions may explain differences of our findings with others, who reported that phage D29, TM4 and DS6A had lytic activity against *Mtb* in acidic conditions (pH 5)(13), and a range of phages had lytic activity against *M. smegmatis* mc^2^155 at pH 5.5 and 6(14).

After adhesion to the host bacterium, phages inject their DNA and use the bacterial replication system for replication and assembly of new phage virions(15). This process precedes lysis of the bacterium in case of a lytic infection. Therefore, a certain degree of bacterial replication is required for phages to have effect, as also shown in the modelling results. In addition, phage virions have been shown to preferentially attach at sites of cell wall synthesis at the poles and septa of mycobacteria and this has been shown for two out of three phages used in this study(16). This can explain our findings of absent lytic activity in models with non-replicating mycobacteria, including stationary phase cultures and cultures in anaerobic conditions. In assays with non-replicating mycobacteria, mycobacterial loads remained stable over time despite incubation with phage. Persistence of phage concentrations in assays with non-replicating mycobacteria, despite phage instability (Supplementary Figure 2), may suggest low grade phage replication in these conditions.

Our findings contrast with previous reports showing activity of a range of phages against *M. smegmatis* mc^2^155 in hypoxia and against stationary phase mycobacteria(14, 17, 18). For *Mtb*, a recent study found that phages D29, TM4 and DS6A had lytic activity in hypoxic conditions in plastic vacutainers(13). Our assay design differs from those in the literature due to the use of phage inactivation buffer before quantification of CFU, to minimize phage carry-over(19). Phage carry-over refers to the unintended persistence of phages in experimental samples or systems after a phage treatment or assay, which can lead to false results or complications in subsequent experimental stages. When culture-based read-outs are used to quantify results of assays with non-replicating mycobacteria this can become an issue due to an increase in effect of phages remaining in experimental samples when mycobacteria start replicating during the culture.

In assays with replicating mycobacteria, there were subtle differences in mycobacterial loads and phage concentrations between phage conditions (Figure 2, Supplementary Table 2-3, Supplementary Figure 6)). The phage combination and Fred had the most pronounced bactericidal activity, whereas their phage concentrations over time were lowest, which was accounted for in the modelling. This is likely due to a predator-prey balance, where phage replication reduces with the reduced availability of bacterial hosts to replicate in(20). Addition of phage was able to prevent regrowth seen in conditions incubated with RIF and INH, despite the occurrence of INH resistance. Our modelling approach can support future study design and possible implementation through simulations of different experimental or clinical scenarios, including different inocula or multiplicities of infection (MOIs), and timing, duration, or sequence of therapies.

Limitations of this study include the sub-optimal control of the anaerobic conditions; future studies may use more robust hypoxia inducing systems like automated anaerobic cultures systems. We used isoniazid activity as a marker of achieving true hypoxia, as its activity is strongly reduced in hypoxic conditions(21). In our anaerobic assay, although not completely absent, isoniazid activity was clearly reduced compared to what was seen in planktonic conditions (proportional change in AUC compared to FAS control -31% in hypoxia versus -49% in time kill assay). Secondly, the use of a culture-based read-out for quantification of non-replicating mycobacteria has limitations, as mycobacteria may remain viable but not culturable(22). In this study we focussed on *Mtb* strain H37Rv, but it would be worthwhile to expand these evaluations of physiologic conditions across multiple lineages(9). There was significant inter-experiment variation of phage stability in acidic conditions, limiting conclusions on lytic phage activity at pH 5.5. A limitation of the model analysis was the minor misprediction of the full trajectory of the combination of RIF with phage combination (Supplementary Figure 6-7), based on the assumption of correct prediction of, and no interaction between, efficacy of RIF and the phage combination. The phage combination was modelled as a single entity but given the even representation of phages in the combination this simplifying assumption was warranted (Supplementary Figure 1-2). RIF efficacy was incorporated into the model without distinguishing between activity against replicating or stationary bacteria, so interpretation of those treatment groups should be taken with care. Only RIF was included as antibiotic in the current modelling framework, however the model allows for expansion by inclusion of other antibiotics.

In conclusion, the phages evaluated here need actively replicating mycobacteria to have lytic activity against *Mtb*. Their activity is reduced in low pH, low oxygen and low growth rate conditions. Further clinical development of phage therapy for TB including phages will need to include regimens with agents that have sterilizing activity against non-replicating mycobacteria, and we suggest that phages should be integrated in early stages of TB regimens when mycobacteria are still actively replicating. Furthermore, here we present a series of tests and analysis methodologies to be used for future evaluations and selection of phages in the therapeutic pipeline for TB.

## Methods

Most experiments were performed at Radboud university medical center (Rumc, Nijmegen, the Netherlands) with the hypoxia assay performed in parallel at Seattle Children’s Research Institute (SCRI, Seattle, USA).

### Bacterial strains and growth conditions

*Mtb* H37Rv (Rumc: ATCC, Manassas, Virginia, USA and SCRI: BEI Resources, USA) was used for all experimental assays. Bacteria were cultured from frozen stocks in 7H9 broth (BD, Maryland, USA) supplemented with 10% oleic acid, albumin, dextrose and catalase (OADC) (BD, Maryland, USA) and 0.05% Tween 80 (Sigma-Aldrich,USA) (pH 6.6) at 37°C with shaking or rotating. For time-kill experiments in planktonic growth conditions and hypoxia, *Mtb* H37Rv was cultured for five days to exponential phase. For experiments with acidic media *Mtb* H37Rv was cultured for seven days to exponential phase in the same medium with the pH decreased by addition of hydrochloric acid (Sigma Aaldrich, USA) to pH 5.5. Single cell solutions were made by passing through a needle and an inoculum of 10^6^ colony forming units (CFU)/milliliter (mL) was obtained by diluting an 0.5 McFarland standard in 7H9 with 10% OADC and 1mM Calcium Chloride (CaCl₂, Sigma-Aldrich, USA). For hypoxic conditions, 96-well plates were prepared containing the exponential phase inoculum at day zero, added to an anaerobic box (A-Box, Kentron Microbiology, Doetinchem, the Netherlands or Thermo Scientific™ AnaeroPack™ 7.0L Rectangular Jar, USA) with anaerobic gas generator sachets (AnaeroGen bags Thermo Scientific, Hampshire, England, or Thermo Scientific™ AnaeroPack™, USA) and test strips (BD, Maryland, USA) for four days to induce a hypoxic state prior to addition of respective treatment conditions (Figure 1). To assess effective phage infection in hypoxia, growth control and phage combination conditions for one experiment underwent a phage inactivation step at Day 31 to inactivate any extracellular phage, whereafter the 96-well plates were incubated for another 14 days in aerobic conditions, before they were sampled for mycobacterial loads. For stationary growth experiments, *Mtb* H37Rv was cultured for 14-21 days and the stock was used undiluted, after passing through a needle, without addition of fresh medium.

*M. smegmatis* mc^2^155 (donated by Edith Houben, Amsterdam University Medical Center, the Netherlands) was used for phage propagation and quantification in soft agar overlays. *M. smegmatis* mc^2^155 was transferred into 7H9 medium supplemented with 10% OADC and 1mM CaCl₂ prior to use for soft agar overlays. For measurement of phage concentrations of different phages in the combination condition, mutant strains of *M. smegmatis* mc^2^155 with differential susceptibility to the respective phages were used (Supplementary Methods, Supplementary Figure 1). *M. smegmatis* mc^2^155 mutant strains IF01-IF04 were cultured in presence of a range of antibiotics (hygromycin, streptomycin, kanamycin and/or anhydrotetracycline).

Whole genome sequencing was performed on isolates incubated with rifampicin (RIF) or isoniazid (INH) for 31 days. Colonies on 7H11 medium were swabbed and resuspended in glycerol broth and supplemented 7H9(23). Glycerol broth samples were heat-inactivated for DNA isolation and sequencing as described before(23). The WHO catalogue was used for calling of mutations associated with resistance(12). Phenotypic susceptibility testing of isolates incubated with INH was done using the broth microdilution assay according to the EUCAST protocol(24).

### Mycobacteriophages

Fred313cpm-1Δ33 (Fred) is a Subcluster A3 phage from which the integrase gene is deleted from Fred313cpm-1, a naturally occurring lytic variant of phage Fred313(9). FionnbharthΔ45Δ47 (Fionnbharth) is a Subcluster K4 phage from which both the repressor and integrase genes are deleted(9). Muddy HRM N0157-2 (Muddy) is a host range mutant of phage Muddy, a naturally lytic Cluster AB phage, with activity against *Mtb*(9). All phages were originally isolated on *M. smegmatis* mc^2^155 in the context of the PHIRE and SEA-PHAGES teaching programmes(25). Phages were propagated on lawns of *M. smegmatis* mc^2^155. For Fionnbharth and Muddy, high-titer lysates were prepared with polyethylene glycol precipitation, and purified with ultrafiltration as described before(26). Fred was further amplified by harvesting high-titer lysates from agar overlays, due to the low initial titer of the received stock, and subsequently passed through a 0.22 µm filter.

### Experimental conditions

The general experimental set-up is demonstrated in Figure 1. On day 0, 96-well plates (one for each sampling day) were inoculated with 100 µl of *M. tuberculosis* achieving a final concentration of 1×10^6^ CFU/mL in each well. A ferrous ammonium sulphate (FAS, Sigma Aldrich, Darmstadt, Germany) treated growth control condition was included in all experiments as FAS could potentially affect viability of *Mtb*. For phage treatment with single phages and the phage combination, phage master mixes were prepared, in 7H9 broth with 10% OADC and 1mM CaCl₂ (pH 6.6; with pH 5.5 for acidic conditions), based on titer and to reach a multiplicity of infection (MOI) of 10:1 (phage versus mycobacteria). 100 µl of phage master mix was added to each well containing bacteria, achieving a total volume of 200 µl per well. Similarly, for RIF (R3501, Sigma-Aldrich, USA) and INH (I3377-5G, Sigma-Aldrich, USA; hypoxia and time-kill experiments only) conditions (both dissolved in dimethyl sulfoxide (Sigma Aldrich, Darmstadt, Germany), 100 µl of antibiotic solution was added to each well to reach a final concentration of 0.1 µg/mL. For stationary growth experiments, the phages and antibiotics were added concentrated in 10 µL of phage buffer (10 mM Tris pH 7.5, 10 mM MgSO4, 68.5 mM NaCl, and 1 mM CaCl2). Phage buffer (without phage or antibiotics) was added to control wells. In a parallel time-kill experiment, the same bacterial stock was diluted in 7H9 broth with 10% OADC and 1mM CaCl₂ and incubated with RIF or the phage combination. Biological triplicates were performed for all conditions. Plates were incubated static at 37°C.

For phage stability testing, phages were incubated in 7H9 broth with 10% OADC and 1mM CaCl₂ (pH 6.6; with pH 5.5 for acidic conditions) and incubated at 4℃ or 37℃.

### Phage and bacterial quantification

For assays with replicating mycobacteria, samples were taken on days zero, three, seven, 14, and 31 (and ten and 21 for time-kill assays) for colony-forming unit (CFU) and plaque-forming unit (PFU) counts. For stationary phase, sampling was performed on days zero, three, seven and 14. For hypoxia, sampling was performed on days zero, four, seven and 14, with day 31 in one single experiment. Supernatants were collected and serial dilutions made in phage buffer were spotted on *M. smegmati*s-containing top agars on 7H10 plates (BD, Maryland, USA) containing dextrose and CaCl₂ for phage counts. Open-source protocols for reagents and top-agars were followed(27). Bacterial pellets were resuspended in FAS for phage inactivation for 5 minutes, spun down (3500 rpm for 5 minutes), and subsequently resuspended in 100 µL 7H9 containing 0.05% Tween 80. For CFU determination on 7H11 plates (Sigma Aldrich, Darmstadt, Germany) containing 10% OADC, 10 μL droplets were spotted in triplicate of 6 serial dilutions and incubated at 37°C for 14-21 days.

### Qualitative assessment of phage activity in hypoxia

In a subset of experiments a high throughput stamping methodology was used to evaluate kinetic effects of phage on *Mtb* in normoxia or hypoxia. The protocol as described above was followed with some minor changes due to laboratory set-up, detailed in the Supplementary Methods.

### Statistical analysis

All experiments were performed in biological and technical triplicates. Results of two independent subsequent experiments were merged for the analysis, unless stated otherwise. The mean ± standard errors are shown in the Figures. R Studio(28) and GraphPad Version 8 (San Diego, CA, USA) were used for statistical analyses and creation of graphs. Areas under the curve (AUC) were calculated for each condition and compared using one-way ANOVA with Dunnett’s test; the FAS control condition was taken as reference due to a decrease in CFU in the FAS condition observed in some experiments. Phage concentrations were compared to the concentration of the phage combination condition. Proportional change in AUC was calculated by division of the mean AUC of an experimental condition of interest by the reference (FAS for CFU, phage combination for PFU).

Mixed-effect modelling and two-way ANOVA were used for comparison between conditions at respective time-points, with Tukey or Dunnett’s test for multiple comparisons, respectively.

### Model-based analysis

To characterize the longitudinal dynamics of the combination of phages and RIF, mechanistic mathematical models were developed, reflecting phage-bacterium interaction dynamics and antibiotic efficacy(29–31). Numerical and graphical diagnostics included parameter values and precision, and simulation-based visual predictive checks(32). Statistical improvement of candidate models for describing phage activity, antibiotic activity, and the combination thereof in comparison to control conditions, was tested by likelihood ratio test of the objective function value, assuming a Χ^2^-distribution (Supplementary Methods).

## Acknowledgements

This project was funded through a European Union’s Horizon 2020 - Marie Sklodowska-Curie Personal Fellowships (grant agreement No 101063247 (Phage-TB); SJ); and the Gates Foundation, INV-060943. This work was also supported by grants to GFH from NIH GM116884, the Howard Hughes Medical Institute GT12053, the Seattle Tuberculosis Research Advancement Center (SEA-TRAC) at Seattle Children’s Research Institute from the National Institute of Allergy and Infectious Diseases funded program under award number P30AI168034, and under R21 grant AI156807-01 (to R.N.C. and S.E.L.). The following reagent was obtained through BEI Resources, NIAID, NIH: *Mycobacterium tuberculosis*, Strain H37Rv, NR-123. We thank Khisi Mdluli (Gates Foundation) for his thoughtful input on experimental set-up.

## Author contributions

SJ, SEL, MPT, CAG, GFH, AHD, RNC and JvI contributed to conceptualization and design of experiments and interpretation of results. Experiments were conducted by SJ, SEL, MPT, TS and IF. SJ, SEL and MPT drafted the work, which was critically reviewed and approved by all co-authors. All authors agree to be accountable for all aspects of the work presented.

## Data sharing statement

The data that support the findings of this study are available from the corresponding author upon reasonable request.

